# Comprehensive analysis of immune evasion in breast cancer by single-cell RNA-seq

**DOI:** 10.1101/368605

**Authors:** Jianhua Yin, Zhisheng Li, Chen Yan, Enhao Fang, Ting Wang, Hanlin Zhou, Weiwei Luo, Qing Zhou, Jingyu Zhang, Jintao Hu, Haoxuan Jin, Lei Wang, Xing Zhao, Jiguang Li, Xiaojuan Qi, Wenbin Zhou, Chen Huang, Chenyang He, Huanming Yang, Karsten Kristiansen, Yong Hou, Shida Zhu, Dongxian Zhou, Ling Wang, Michael Dean, Kui Wu, Hong Hu, Guibo Li

**Author notes:** These authors contributed equally to this work (Jianhua Yin, Zhisheng Li, Chen Yan, Enhao Fang, Ting Wang). These authors contributed equally to this work (Ling Wang, Michael Dean, Kui Wu, Hong Hu, Guibo Li).

## Abstract

The tumor microenvironment is composed of numerous cell types, including tumor, immune and stromal cells. Cancer cells interact with the tumor microenvironment to suppress anticancer immunity. In this study, we molecularly dissected the tumor microenvironment of breast cancer by single-cell RNA-seq. We profiled the breast cancer tumor microenvironment by analyzing the single-cell transcriptomes of 52,163 cells from the tumor tissues of 15 breast cancer patients. The tumor cells and immune cells from individual patients were analyzed simultaneously at the single-cell level. This study explores the diversity of the cell types in the tumor microenvironment and provides information on the mechanisms of escape from clearance by immune cells in breast cancer.

**One Sentence Summary:** Landscape of tumor cells and immune cells in breast cancer by single cell RNA-seq

---

Breast cancer is the most common cancer and the leading cause of death from cancer in women worldwide(1). Four subtypes of breast cancer with distinct expression profiles have been classified based on gene expression signatures associated with highly variable clinical characteristics(2, 3). To design targeted treatment for such a diverse disease, understanding its molecular mechanism of initiation and progression is essential(4). Several studies have shown that the presence of tumor-infiltrating lymphocytes (TILs) is associated with breast cancer progression and neoadjuvant chemotherapy response(5-7). TIL levels within and between different subtypes of breast cancer vary(8). Based on the CD8^+^ T cell infiltration phenotype, tumors can be categorized as T-cell-inflamed and non-T-cell-inflamed tumors(9). In T-cell-inflamed tumors, the tumor cells act together with the tumor microenvironment (TME) to inhibit the antitumor functions of T cells. This process results in an exhausted T cell phenotype. Non-T-cell-inflamed tumors escape immune system clearance by preventing T cell infiltration into the TME(10). Cancer-associated fibroblasts (CAFs), tumor-associated macrophages (TAMs) and oncogenic pathway alterations in tumor cells are reportedly responsible for the non-T-cell-inflamed phenotype(10-14). However, the mechanism of immune evasion in breast cancer remains unclear. A recent study showed that a high number of TILs was beneficial for triple negative breast cancer (TNBC) and HER-2-positive breast cancer patient survival, but this parameter was an adverse prognostic factor for luminal HER2-negative breast cancer patient survival(15). High-resolution mapping of the TME composition and cell states of different breast cancer subtypes will help us understand the different effects of TILs and the mechanism of tumor immune evasion in breast cancer.

In the past five years, the development of high-throughput single-cell RNA sequencing has enabled high-resolution studies of biological processes(16). Single-cell transcriptome profiling of tumor cells has been used to characterize heterogeneous tumor cells and tumor-associated stromal and immune cells (17-19). To understand the mechanisms of escape from clearance by immune cells in breast cancer further, we molecularly dissected the breast cancer TME by using single-cell RNA-seq.

We performed single-cell RNA-seq on cells isolated from the tumor tissues of 15 human breast cancer patients (Table S1). Our samples included four subtypes of breast cancer: luminal A (P01, P02, P03, P04, P05, P06, P07A, P07B and P08), luminal B (P09 and P10), HER2+ (P11 and P12) and TNBC (P13, P14 and P15). We isolated single cells from tumor tissues without surface marker preselection. With proper quality control and filtration, we obtained single-cell transcriptome data for 52,163 individual cells (Fig. 1A).

**Fig. 1.**
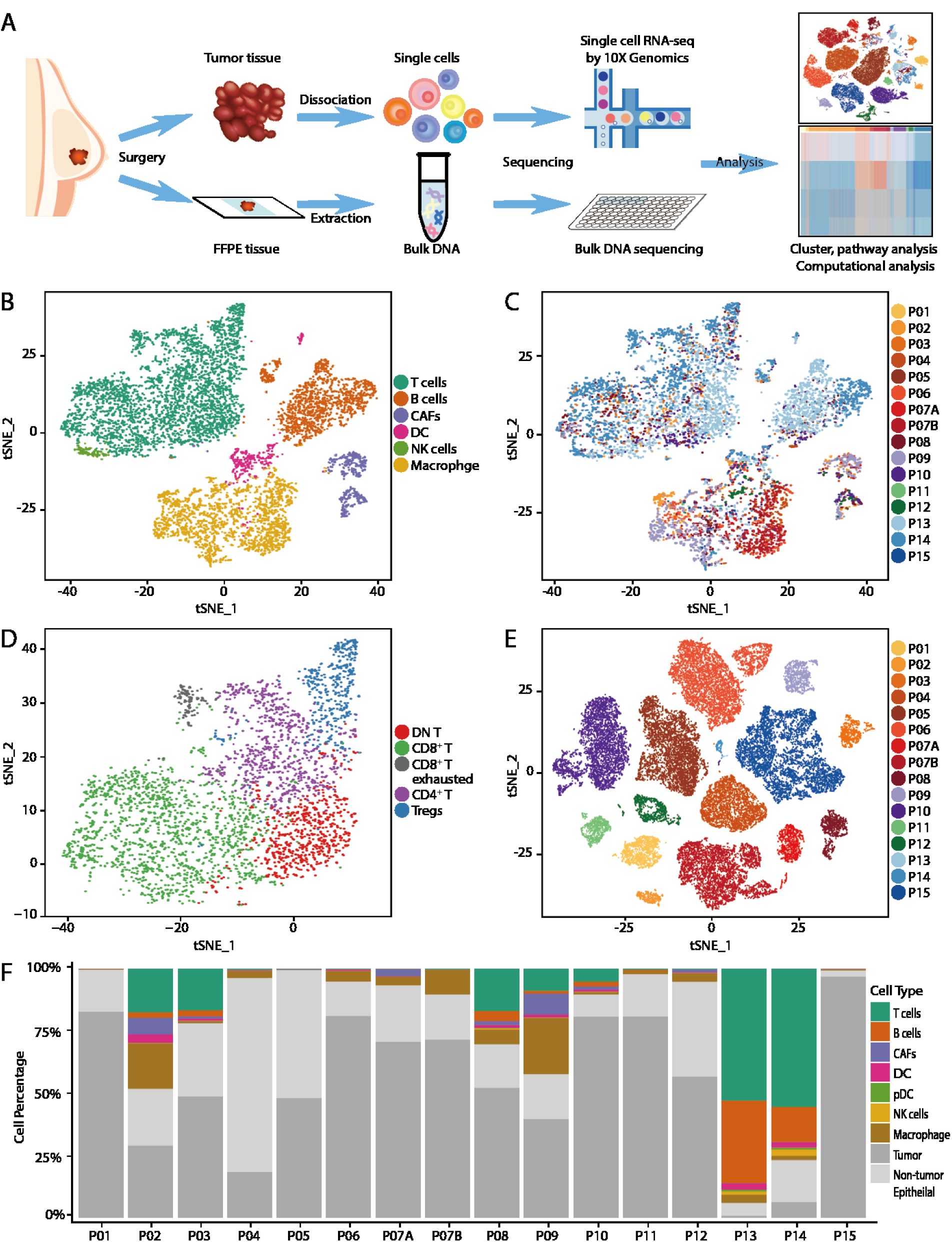
Landscape of TME in breast cancer according to single-cell RNA-seq. (A) Overview of study design. (B and C) t-SNE plot of immune cells and CAFs. The cells are annotated as different cell types according to differentially expressed cell markers. Left: The cells are colored according to the identified cell types (B). Right: The cells are colored according to the sample type (C). (D) t-SNE plot of T cells. T cells were identified as DN T, CD4^+^ T, CD8^+^ T, CD8^+^ exhausted T and Tregs based on specific cell markers and gene signatures. (E) t-SNE plot of tumor cells across 16 samples. The cells are colored according to the sample type. (F) Proportion of different cell types in tumor tissues across the patients.

To analyze the composition of the breast cancer TME, we clustered the cells according to their expression profiles. Preliminary clustering was applied to all cells using the Seurat package (Fig. S2, A, B and C). Each cluster was designated as epithelial or non-epithelial, and the epithelial cells were distinguished further as tumor cells or non-tumor epithelial cells based on the inferred copy number variation information (Fig. S3). The clusters of non-epithelial cells were annotated as T cells, macrophages, B cells, dendritic cells, NK cells and CAFs (Fig. 1, B and C). Based on specific cell markers, the T cell clusters were annotated further as CD4^+^ T cells, regulatory T cells (Tregs), CD8^+^ T cells, and CD8^+^ exhausted T cells (Fig. 1D). The precise identification of T cells will assist in analyzing the immune evasion mechanism of each sample. The tumor cells from different patients clustered separately, suggesting a high degree of inter-tumor heterogeneity (Fig. 1E). Finally, the cell type percentages of individual patients were calculated (Fig. 1F). The concentration and composition of the TILs were variable between different patients, which revealed the complexity of the tumor tissues. In the P13 and P14 samples, which were both TNBC and had a high number of TILs, most of the TILs were T cells and B cells. In many of the luminal A and luminal B samples (P02, P04, P06, P07A, P07B, P08 and P09), macrophages constituted the first or second largest TIL population.

Gene expression profiling and massively parallel sequencing have identified four main molecular subtypes of breast cancer; these subtypes have significant genetic and phenotypic diversity and inter-tumor heterogeneity among patients(2, 3). To further analyze the tumor cell heterogeneity within a patient, we examined the breast cancer subtype and oncogenic pathways based on our single-cell RNA-seq data. Unsupervised clustering was applied to each sample, and each subpopulation had a distinct expression pattern (Fig. 2A and 2B). Differentially expressed genes were analyzed for each subpopulation, and their functional characteristics were classified (Fig. 2C and 2D).

**Fig. 2.**
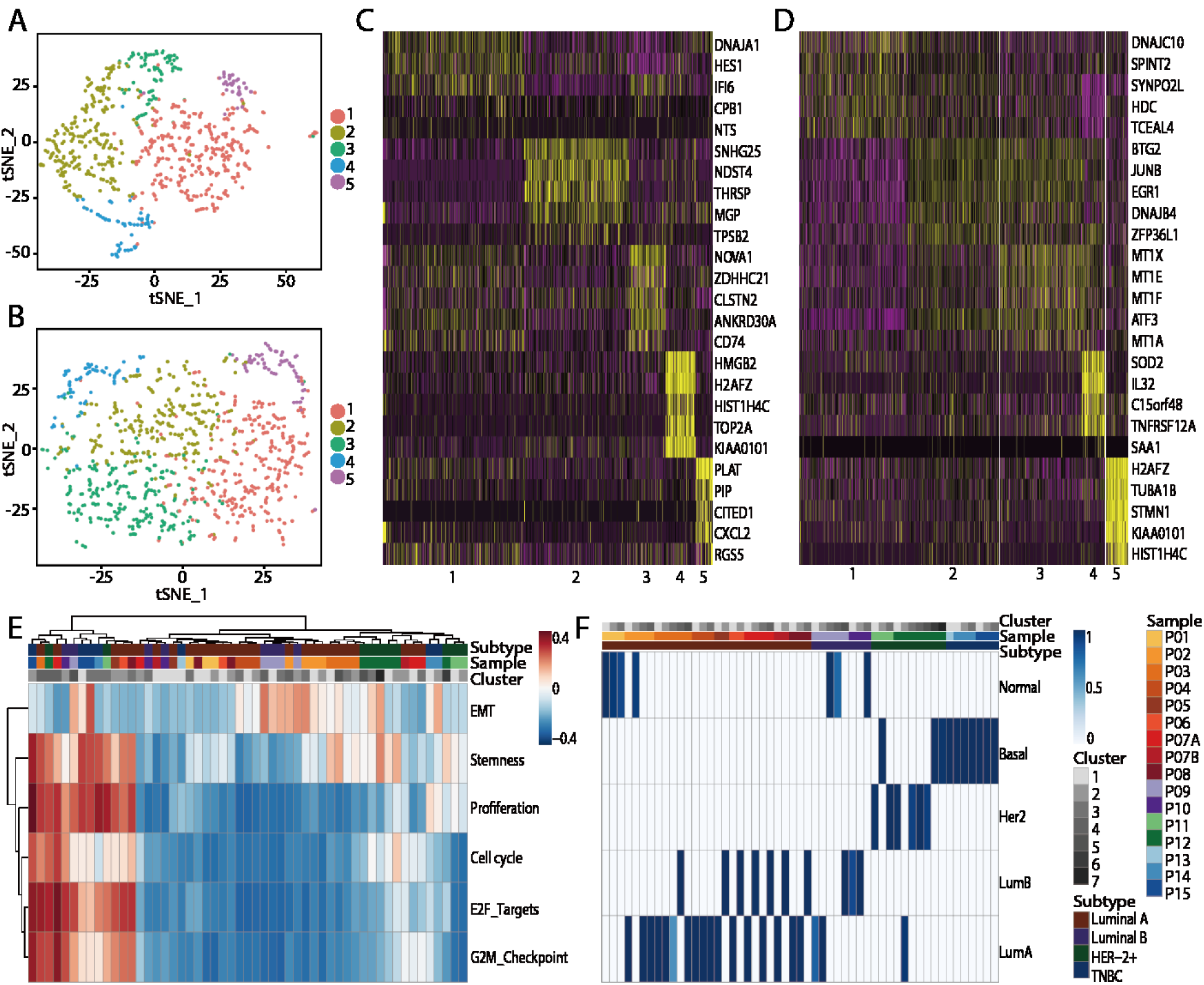
Tumor cell heterogeneity in breast cancer. (A) t-SNE plot of tumor cells in sample P03. The cells are colored according to their clusters. (B) t-SNE plot of tumor cells in sample P09. The cells are colored according to their clusters. (C). Heatmap based on genes differentially expressed in the five clusters of sample P03. The clusters are indicated by color. (D) Heatmap based on genes differentially expressed in the five clusters of sample P09. The clusters are indicated by color. (E) GSVA of genes signatures associated with EMT and cell cycling in each cluster of tumor cells across the patients. (F) Molecular subtype classified by PAM50 in each cluster of tumor cells across the patients.

We also used gene set variation analysis (GSVA) to identify the diversity of cancer phenotypes and signaling pathways among patients(20). EMT programs in cancer have been widely considered as potential triggers of drug resistance, invasion, and metastasis; however, using EMT data for cancer diagnosis and treatment is difficult because their patterns and significance in human epithelial tumors in vivo are unclear(21). To understand EMT patterns in breast cancer in vivo further, we analyzed the EMT-associated gene signatures of the patients. Our results showed that the EMT programs displayed both inter-tumor and intra-tumor heterogeneity. Different subpopulations in the same patient exhibited distinct patterns of EMT programming, which suggest that intermediate states of the EMT program exist in vivo (Fig. 2E). An analysis of gene signatures associated with proliferation and cell cycle identified proliferating clusters in several samples. In sample P15, almost all clusters displayed a high proliferation gene signature score, suggesting that most of the cells in this sample were in active proliferation states. Only a few TILs were detected in sample P15, suggesting that this cancer progressed rapidly without an antitumor immune response. We also analyzed the activation of a series of oncogenic pathways for each individual cluster in each sample, many of which were diverse across the clusters; these data provided us with the resources to understand tumor cell heterogeneity (Fig. S4).

We used the PAM50 classifier to predict breast cancer subtype (luminal A, luminal B, Her2, basal and normal-like) for individual tumor cell clusters in each sample(22). Intra-sample heterogeneity was present in our results showing that the subpopulations of one sample corresponded to different breast cancer subtypes (Fig. 2F). Six luminal A samples (P03, P05, P06, P7A, P7B and P08) also contained clusters conforming to the luminal B subtype. Our results have revealed intra-tumor heterogeneity in breast cancer that could not be detected by bulk RNA-seq or IHC staining.

Our scRNA-seq data highlight the immune cell and CAFs diversity in the breast cancer TME. We identified 9 clusters of T cells, 5 clusters of TAMs, 5 clusters of B cells, 2 clusters of DCs and 2 clusters of CAFs in our samples, and each of these clusters has a distinct expression pattern (Fig. S5A and S5B). To examine the functional state of these cell clusters, we performed GSVA to identify functional phenotype diversity in immune cells and CAFs.

Among the 9 clusters of T cells, 5 of them were annotated as CD8^+^ T cells, and 3 of them were annotated as CD4^+^ T cells. T cell-mediated cytotoxicity is critical for tumor cell clearance. Based on the functional T cell associated gene signature analysis, the 5 clusters of CD8^+^ T cells gradually activated cytotoxic and exhausted T cell gene signatures (Fig. 3A)(23). The clusters annotated as CD8^+^ T_exhausted_ and CD8^+^ T-4 expressed similar levels of cytotoxic genes, but multiple inhibitory receptors were overexpressed in the exhausted T cell cluster; this overexpression may attenuate the cytotoxicity of this T cell cluster (Fig. 4C). Thus, among these five CD8^+^ T cell clusters, T cells in the CD8^+^ T-3 and CD8^+^ T-4 clusters may be able to kill tumor cells. The three CD4^+^ T cell clusters also had varied T cell functional states. One cluster was annotated as regulatory T cells, naïve CD4^+^ T-2s and CD4^+^ T-1s with an exhausted signature (Fig. 4B and 4C). T cells in these three CD4^+^ T cell clusters might regulate CD8^+^ T cell function to different degrees.

**Fig. 3.**
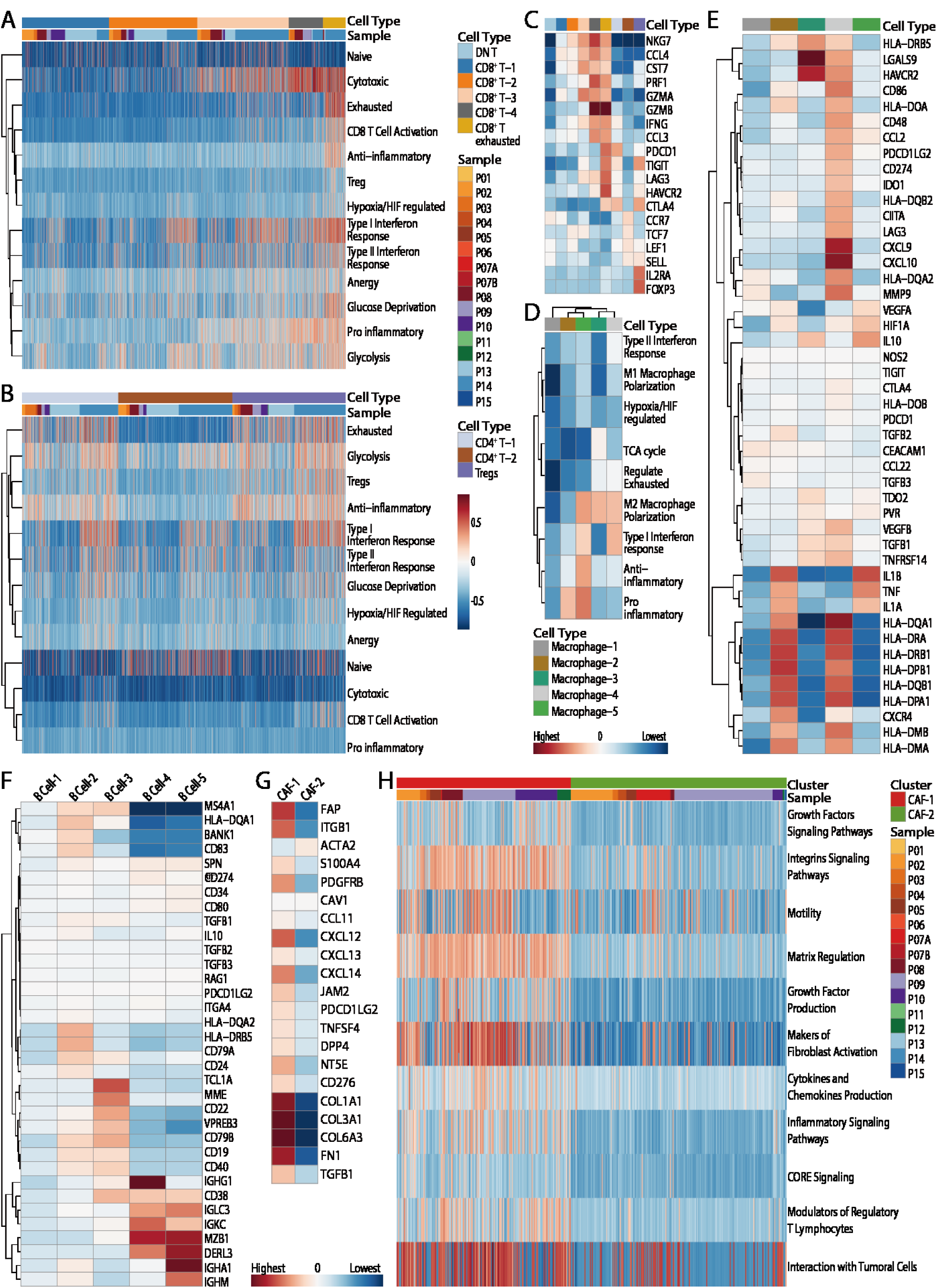
Diversity of immune cells and CAFs in breast cancer. (A) Heatmap showing GSVA scores for gene signatures associated with T cell functions. (B) Heatmap showing the average expression of T cell functional genes in each cluster of T cells: cytotoxic (NKG7, CCL4, CST7, PRF1, GZMA, GZMB, IFNG and CCL3), exhausted (PDCD1, TIGIT, LAG3, HAVCR2 and CTLA4), naïve (CCR7, TCF7, LEF1 and SELL), and Tregs (IL2RA and FOXP3). (C) Heatmap showing GSVA scores for the gene signatures associated with macrophage functions. (D) Heatmap showing the average expression of macrophage functional genes in each cluster of macrophages. (E) Heatmap showing the average expression of B cell functional genes in each cluster of B cells. (F) Heatmap showing the average expression of CAF functional genes in each cluster of CAFs. (G) Heatmap showing the GSVA scores for gene signatures associated with CAF functions.

**Figure 4.**
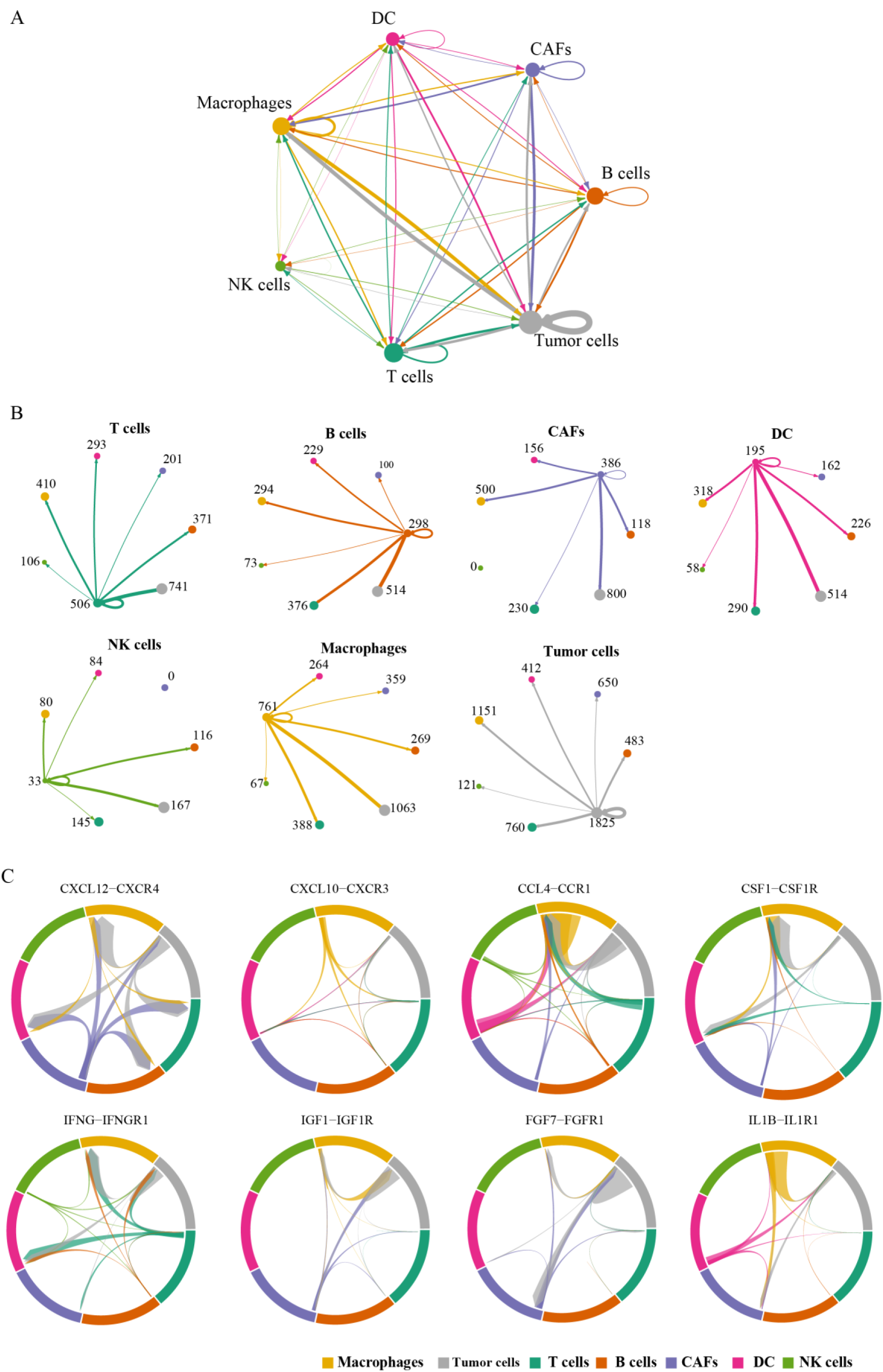
A network of intercellular communication in breast cancer TME. (A)Network of potential interactions between different cell types in breast cancer TME. (B) Detailed numbers of ligand-receptor pairs between indicated cell types. (C)Selected ligand-receptor pairs in breast cancer TME.

TAMs, a major type of leukocytes that infiltrate tumors, play an important role in tumor growth and progression(24). Macrophages in different TMEs are functionally distinct. Our single-cell RNA-seq data identified 5 clusters of macrophages, revealing a high complexity of TAMs in vivo. Clusters 3, 4 and 5 manifested mainly in the M2 state, which may promote tumor growth by suppressing antitumor immune responses (Fig. 3D). We also examined the expression of a series of genes associated with the immune regulatory function of macrophages in these clusters (Fig. 3E). Macrophages in clusters 2 and 4 expressed higher levels of HLA-II genes than the other 3 clusters. In cluster 4, we also detected the expression of CD274, PDCD1LG2 and IOO1, which have been reported to be associated with suppressing CD8^+^ T cell functions.

We also analyzed the expression pattern of several B cell markers, which were previously reported to be associated with B cell maturation and regulatory functions in five B cell clusters (Fig. 3F). Cluster 2 was HLA-II-positive and may be able to present antigens. Cluster 3 was TCL1A-positive; TCL1A-positive cells are reportedly involved in controlling cervical cancer development(25). B cells in clusters 4 and 5 expressed high levels of immunoglobulin-associated genes, suggesting that they were antibody secretion B cells, which might be involved in tumor immune regulation(26).

A recent study had identified four distinct CAF subsets in breast cancer; based on this classification, we examined the properties of CAFs in our data(27)(Fig. 3F and 3G). CAF-1 and CAF-2 showed distinct expression patterns of CAF markers and pathways. CAF-1 cells expressed higher levels of immune suppression associated genes. This finding suggests that CAF-1 cells may be associated with tumor immune evasion in breast cancer.

The cross-talk between tumor cells and stromal cells in tumor microenvironment is critical for cancer initiation and progression. Based on our single cell RNA-seq data, we analyzed intercellular communication in breast cancer TME using a dataset of human ligand receptor pairs(28). A network of potential cell-cell interactions was constructed showing extensive communications between tumor cells and stromal cells (Fig. 4A). Macrophages had the most potential connections with tumor cells (Fig. 4B). The interactions between macrophages and tumor cells mainly mediated by multiple ligand-receptor pairs, including CSF1-CSF1R, IGF1-IGF1R and IL1B-IL1R1, which may be associated with macrophages stimulation and response (Fig. 4C). CAFs densely communicated with tumor cells and macrophages through CXCL12-CXCR4, IGF1-IGF1R and FGF7-FGFR1(Fig. 4C). Macrophages recruited CXCR3 positive T cells by producing CXCL10 and CXCL9, and CCL4 secreted by T cells and DC may regulate recruitment of CCR1 positive macrophages to TME. Besides interactions with known functions, we also identified several potential intercommunications in breast cancer TME, which need further confirmation in future.

Class I human leukocyte antigen (HLA) is responsible for cancer-specific neoantigen presentation(*29*). Genetic alterations in HLA-I molecules are reportedly associated with the escape of tumor cells from immune cell clearance(30, 31). To analyze the mechanism of immune evasion in breast cancer comprehensively, we first examined HLA-I gene expression in the tumor cells at the single-cell level (Fig.5A). The expression of HLA-associated genes was diverse across patients. In most samples (P01, P03, P04, P05, P06, P07A, P07B, P09, P10, P11, P12, P14 and P15), HLA-I gene expression was positively correlated with the percentage of infiltrating T cells. In one TNBC sample (P15) that lacked lymphocyte infiltration, all of the HLA-I genes were downregulated. In contrast, HLA-I genes were overexpressed in sample P14, and the tumor tissue of this patient was highly T cell-inflamed with 50% infiltrating T cells. These results suggest that in samples P01, P05, and P15, HLA-I gene downregulation played a critical role in preventing immune cell infiltration into the tumor tissues.

Our single-cell analysis of tumor cells also revealed diverse HLA-I expression in different subpopulations of the same sample. In sample P02, only cluster 3 showed HLA-I gene expression, which may be adequate for priming the antitumor immune response. In samples P04, P06 and P12, HLA-I gene expression could be detected, but only a few infiltrating T cells were found, suggesting that another T cell exclusion mechanism was involved in these samples.

In samples P02, P03, P08, P09, P10, P13 and P14, over 10% of infiltrating T cells were detected. In these samples, the antitumor immune response was primed, and T cells were recruited to the tumor tissues but failed to kill all of the tumor cells. The T cell composition of each patient, which was diverse among patients, revealed the functional T cell states of individual patients (Fig. 5B). These results showed that in infiltrating T cell populations, the percentage of cytotoxic CD8^+^ T cells (CD8^+^ T-3 and CD8^+^ T-4) in individual samples was less than 35%, which may be not adequate for killing all of the tumor cells. In addition, the existence of regulatory T cells in the tumor tissues of these patients may also inhibit the antitumor function of cytotoxic CD8^+^ T cells. We also used the GSVA scores of T cells and macrophages to further evaluate the functional T cell and macrophage states in individual samples (Fig. S6B and S7; Fig. 5C and Fig. 5D). Cytotoxic scores revealed the ability of T cells to destroy tumor cells, the exhaustion score showed the dysfunctional state of the T cells, and the presence of Tregs and M2s revealed inhibitory T cells and macrophages, respectively. In sample P09, more than 50% of the TILs were macrophages, and most of the macrophages displayed the M2 phenotype, which could protect the tumor cells from T cell attack. For this sample, macrophages may be targeted for therapy. In sample P14, tumor cells expressed high levels of IDO1, LGALS9, CEACAM1, TNFSF10 and PDCD1LG2, which were all reported to inhibit cytotoxic T cell function (Fig. S6A). In sample P14, 8% of the infiltrating T cells were CD8^+^ exhausted T cells, and 12% were CD4^+^ exhausted T cells, which may directly contact the tumor cells and become exhausted. Immune checkpoint therapy may be effective for this patient. In samples P10 and P13, most of the infiltrating T cells had neither cytotoxic nor exhausted signatures, which suggests that these T cells were not activated in the tumor tissues. Effectively activating the T cells in these samples may be critical for treatment. Taken together, analyzing tumor cells and infiltrating immune cells simultaneously by single-cell RNA-seq enables us to understand the precise status of tumor tissues in individual patients comprehensively. This information may be very helpful for clinical diagnosis and therapy.

**Fig. 5.**
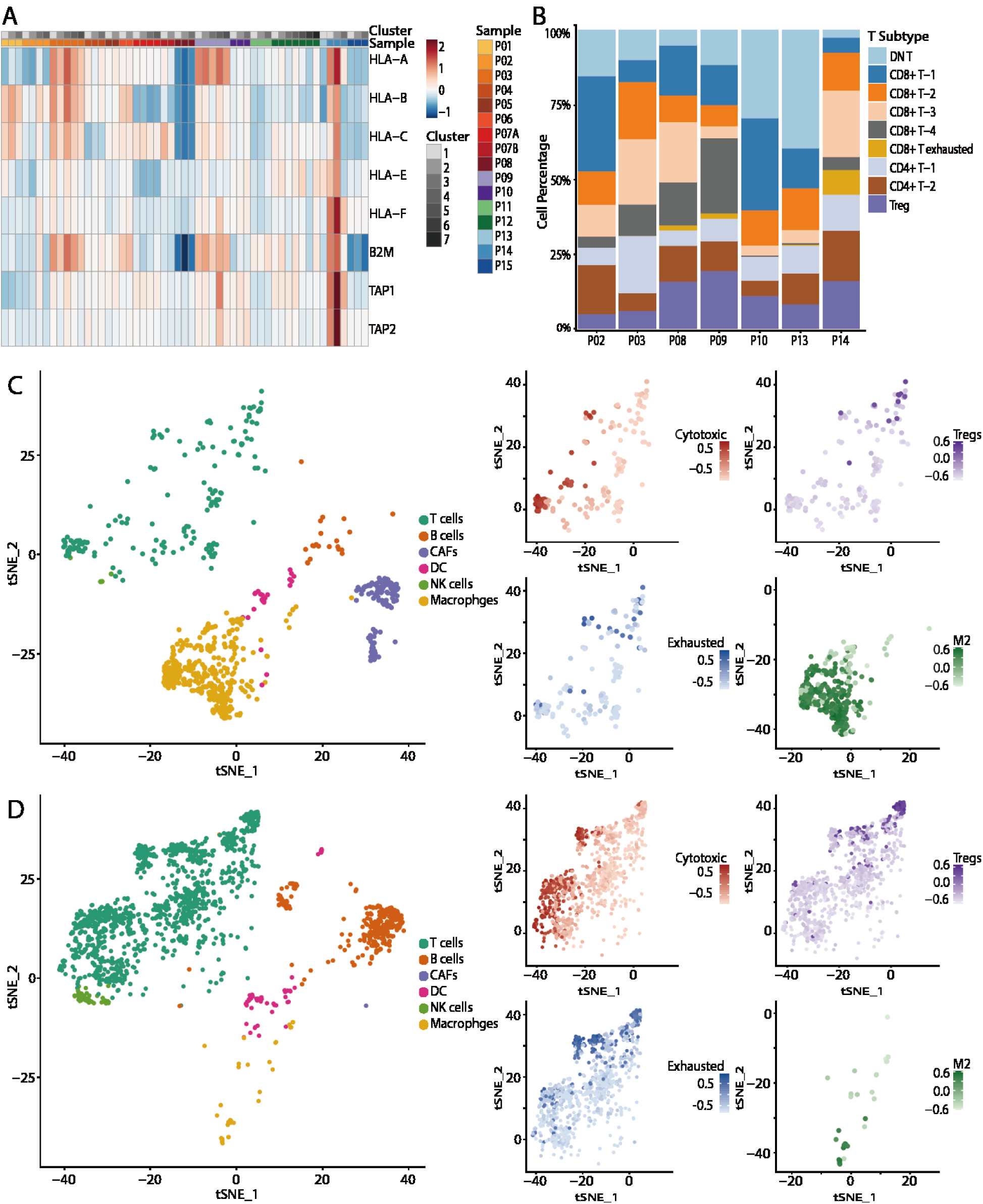
Immune evasion in individual patients. (A) Heatmap showing the average expression of HLA-I-associated genes in each subtype. (B) Composition of T cells in T cell-inflamed patients. (C and D) t-SNE plot of immune cells and CAFs in samples P09 (C) and P14 (D). t-SNE plot of T cells and macrophages showing the T cell functional state (GSVA scores for cytotoxic, Treg and exhausted pathways) and macrophage M2 signatures.

Using single-cell transcriptomic data from 53,000 cells, we dissected precisely the TME of 15 breast cancer patients by determining the proportion of each cell type and their gene signatures. Several subpopulations of immune cells and CAFs were identified. This finding highlights the diversity of macrophages, CAFs and B cells in the breast cancer TME. The mechanism of T cell exclusion and T cell exhaustion in the breast cancer samples was analyzed comprehensively based on the landscape of the TME.

Our results revealed that breast cancer tumor cells appear to interact with TAMs and CAFs to escape immune system clearance. Several signaling pathways and molecular factors that are likely involved in this process were identified, as well as complex and diverse active mechanisms in each sample. Thus, single-cell RNA-seq provides comprehensive insights into the immune suppression network in the TME, and this information could be a critical guide for personally precise therapy design for cancer patients.

## Acknowledgment

This work was supported by the Shenzhen Municipal Government of China (grant GJHZ20170314152701465), National Natural Science Foundation of China (No.31500694, 81672593 and 81272899), Natural Science Foundation of Guangdong Province, China (2017B020227012), “sanming” project of medicine in Shenzhen(SZSM201512015). We also acknowledge in part by the Intramural Research Program of the National Institutes of Health, National Cancer Institute, Center for Cancer Research, and Division of Cancer Epidemiology and Genetics from Leidos-Frederick under contract # HHSN261200800001E.

## Supplementary Materials

### Materials and Methods

#### Patients and tumor specimens

We chose 15 patients with four pathologic subtypes: luminal A (P01-P08), luminal B (P09, P10), HER-2 (P11, P12) and TNBC (P13-P15). These molecular subtypes were diagnosed by pathological examination. Five samples (P01, P03, P05, P06 and P14) were patient tissue biopsies and were diagnosed as breast cancer by highly qualified pathologists; the other samples were surgical samples. Two samples were postoperative samples after neoadjuvant chemotherapy (P12 and P15), and the others were from chemotherapy-naïve patients. This study was approved by the Institutional Review Board of BGI (BGI-IRB, N0: BGI-IRB 17090) and Shenzhen People’s Hospital (Second Clinical Medical College of Jinan University). In addition, all patients who provided specimens signed an informed consent form and agreed to the specimens being used for scientific research. Sample P07 was from two different parts of the tumor, and sample P08 was from a male breast cancer patient. In all, 16 primary tumor specimens were collected and processed for single-cell RNA sequencing.

#### Tumor disaggregation and single cell collection

Fresh tumor tissue samples were minced into tiny cubes <1 mm^3^on ice and transferred into a 1.5-mL tube contenting 20 U/ml collagenase III (Invitrogen), 3 U/ml hyaluronidase (Invitrogen), 1 U/ml DNase I (Invitrogen) and 1× Hank’s balanced salt solution (HBSS, Invitrogen). The tumor pieces were digested in this digestion media for 60 min at 37°C. Then, the tumor cells were filtered by a 40-μm cell strainer (BD) and centrifuged at 300 g and 4°C for 5 min. After centrifugation, the supernatant was discarded, and the cell pellet was re-suspended in PBS with 0.02% BSA (Sigma).

#### Single-cell RNA sequencing

Cells were concentrated to 700-1000 cells/μl and loaded on a Chromium Single-Cell Instrument (10X Genomics) to generate single-cell gel bead-in-emulsions (GEMs). For cDNA recovery, amplification and library construction were performed with a Chromium^™^ Single Cell 3’ Reagent Kit v2 (10X Genomics) according to the manufacturer’s protocol. Sequencing libraries were loaded in BGIS EQ-500 using the following read lengths: 26-bp read 1 (containing the 16-bp 10x^™^ barcode and 10-bp randomer), 100-bp read 2 and 8-bp read 1 barcodes (i7 index).

#### Bulk DNA extraction and sequencing

Genomic DNA from formalin-fixed and paraffin-embedded (FFPE) tumor and adjacent normal tissues was extracted using a QIAamp FFPE DNA kit (QIAGEN) according to the manufacturer’s instructions. Whole genome shotgun (WGS) libraries were constructed using a BGIseq FFPE DNA Library Prep Kit (BGI). Briefly, 20 μl FFPE DNA was fragmented and end-repaired with 10 μL one-step Fragment & End-repairing Buffer (BGI) at 37°C for 20 min and 65°C for 15 min. Then, the indexed adenylated 3’ adapters and 5’ adapters were ligated to the DNA fragments by T4 DNA ligase (NEB), and the ligated products were purified by 0.5× Ampure XP beads (Agencourt). Pre-PCR reactions were performed with 20 μL purified DNA, 25 μL 2× Kapa HiFi supermix (Kapa), and 5 μL PCR primers. The program was as follows: 95°C for 3 min; 9 cycles of 98°C for 20 s, 60°C for 15 s, and 72°C for 40 s; then, 72°C for 5 min. Pre-PCR products were purified with 1.2× AMPure XP beads and eluted in 40 μL nuclease-free water. Subsequently, the eluted DNA was divided into two parts. One was used directly to make single strand circular DNA, which yielded the final WGS library.

The other part was used to construct whole-exome sequencing (WES) libraries, which were first captured by BGIseq Human Exome V4 Kit (BGI) according to the manufacturer’s instructions. For this process, 40 μL captured exon DNA was subjected to post-PCR with 50 μL 2× Kapa HiFi supermix and 10 μL PCR primers. The program was as follows: 95°C for 3 min; 13 cycles of 98°C for 20 s, 60°C for 15 s, and 72°C for 40 s; then, 72°C for 5 min. Lastly, 1.2× AMPure XP bead-purified post-PCR products were used to make single strand circular DNA, which yielded the final WES library. WGS libraries were sequenced at an average coverage of 3∼5× on a BGISEQ-500 sequencer with 100 bp paired-end reads, while WES libraries were sequenced by BGI in China at an average coverage of 300× on a BGISEQ-500 system with 100 bp paired-end reads.

#### WES and Bioinformatics

Genomic DNA was extracted from patient peripheral blood samples. A BGI Exome Enrichment Kit v4.0 was used to enrich the exome capture; then, the captured products were circularized. Rolling circle amplification (RCA) was performed to produce DNA nanoballs (DNBs). Each resulting captured library was then loaded on a BGISEQ-500 sequencing platform and was sequenced according to the manufacturer’s instructions. Sequencing-derived raw image files were processed by BGI Seq500 base-calling software with default parameters, and the sequence data for each individual were generated as 100-bp paired-end reads, which were defined as "raw data" and stored in the FASTQ format. Then, analysis workflows were used to perform sequencing read alignment, variant calling, and variant annotation. After filtering the raw data, all clean data from each sample were mapped to the human reference genome (GRCh38/HG38). Burrows-Wheeler Aligner (BWA) software was used to perform the alignment(32). Variants were identified by the Genome Analysis Toolkit according to the recommended best practices (GATK, https://www.broadinstitute.org/gatk/guide/best-practices). The sequencing depth and coverage for each individual were calculated based on the alignments. The hard-filtering method was applied to achieve high-confidence variant calls. Then, the SnpEff tool (http://snpeff.sourceforge.net/SnpEff_manual.html) was applied to perform a series of annotations for the variants. The final variants and annotation results were used in the downstream advanced analysis. The variant frequency and function prediction scores were annotated in public databases, including dbSNP, the 1000 Genomes Project, ExAC and ESP, as well as an in-house BGI database and prediction software, such as SIFT, PolyPhen, MutationTaster and GERP for non-pathogenic variant filtration(33-36).

#### Single-cell RNA sequencing data processing

Sequencing data from the BGI Seq500 sequencer were paired-end reads that were used for sequencing library construction with the GemCode Single-Cell 3’ Library V1 Kit; the first read consisted of 14-bp cell barcodes and 10-bp unique molecular identifiers (UMIs), and the second read consisted of 100-bp 3’ transcripts and 8-bp sample indexes. For sequencing library construction using the GemCode Single-Cell 3’ Library V2 Kit, the first read consisted of 16-bp cell barcodes and 10-bp UMIs, and the second read consisted of 100-bp 3’ transcripts and 8-bp sample indexes. Then, we used CellRanger v1.2 and v2.0 software to process the raw FASTQ files, align the sequencing reads to the GRCh38 reference transcriptome build using STAR and generate a filtered UMI expression profile for each cell (37, 38).

#### Preliminary filtration and clustering

Data from all samples were merged into one gene cell barcode matrix. This gene cell barcode matrix was filtered based on the gene number per cell (only cells with 400 to 8000 genes were kept) and mitochondrial UMI count proportions (only cells with less than 20% of mitochondrial UMI counts were kept). Genes that were present in at least 10 cells in one sample were kept for further analysis. In total, 52614 single cells and 23383 genes passed the QC criteria. The Seurat package was used to analyze our single-cell data(39). Highly variable genes were calculated with the Find Variable Genes method of the Seurat package; these genes had a mean expression value between 0.0125 and 4 and a dispersion greater than 0.5. The selected genes were used for principal components analysis (PCA) dimension reduction on a log-normalized data matrix. The cells were clustered into 40 groups by the FindClusters method using the first 45 principal components, which were further assigned as epithelial, non-epithelial and unclassified.

#### Copy number variation (CNV) inferring and tumor identification

To select the tumor cells from the normal epithelial cells, we chose the inferCNV package to calculate the CNV score for each cell(40). A peripheral blood mononuclear cell (PBMC) sample from a healthy donor that was sequenced on BGI Seq500 served as the reference for inferCNV. Such data were filtered so that cells with less than 200 genes, more than 2000 genes, more than 5% mitochondrial UMI counts, or more than 10000 UMI counts were filtered. Both the reference data and our sample data were transformed into log2(TPM+1), where TPM (transcripts-per-million) was computed as the proportion of UMIs of a gene within each cell multiplied by 1,000,000. Genes that were less than 4.5 in log2((TPM+1)/10) space were removed. CNV scores were computed on a moving average window equal to 101. Scores were bounded to −1 to 1, and any score between - 0.2 and 0.2 would be set to 0. Instead of using the hierarchical clustering that inferCNV provided, we performed k-means clustering (k=2) on the CNV scores of the reference data and sample data. The cluster with a center that had a lower sum of squares than the CNV score, which contained most of the reference cells, was considered normal, while the other scores were considered tumors. To examine the validity of the CNV score, we applied the following transformation to the WES data. First, the ratios of the WES data from tumor tissue and normal tissue from the same patient were calculated and normalized by their average depth in log space. The ratio data were segmented and smoothed by a DNA copy package. The output was also bounded to −1 to 1. Finally, the segmented ratio data were mapped to the gene list according to their genomic position. The results showed that the single-cell inferred CNV scores matched the WES CNV pattern (Fig. S2).

#### Non-epithelial cell classification

The 7804 cells that were categorized as non-epithelial cells were combined, and PCA was repeated on a list of 344 selected genes. These cells were clustered into 24 groups by the Seurat FindClusters method using the first 20 principal components with resolution=2, k.param=20 and k.scale=15. Cluster-specific genes were used to identify the cell types. T cell clusters were indicated by CD3D, CD2, CD3E and CD3G enrichment. B cells were distinguished by CD19, CD79A and CD79B enrichment. NK cells were annotated based on the enrichment of the markers NKG7 and GNLY. Macrophages were annotated based on the enrichment of the markers CD163, CSF1R and CD14. T cell clusters were further classified as CD4+ T cells, CD8+ T cells, double-negative T cells, CD8+ exhausted cells and regulatory T cells. To show differences in genes expression between the clusters, AverageExpression in Seurat was performed on the log scaled data.

#### GSVA

We applied GSVA to identify the molecular phenotypes of single cells using the log_2_(TPM?+?1) data(20). First, we obtained GSVA scores for 39 oncogenic gene sets (MSiDB version 6.1) for each tumor cell(41). Thirteen gene sets were found for the T cells, 9 gene sets were found for the macrophages, and 11 gene sets were found for the CAFs. Then, the GSVA scores of the cells for each cluster from each sample were averaged, and the values were plotted as a heatmap using R package “pheatmap” (https://cran.r-project.org/web/packages/pheatmap/index.html).

#### Breast cancer subtype prediction

We utilized the PAM50 classifier to predict the intrinsic breast cancer subtype (normal-like, basal, Her2, luminal B, and luminal A) for each tumor cell cluster of each sample(22). We averaged the gene expression value of 50 genes of the cells for each tumor cluster from each sample for PAM50. Then, we used the average gene expression values of each tumor cell cluster to plot a heatmap, and we used the confidence score of each subtype calling according to the PAM50 classifier for each tumor cell cluster to plot a heatmap.

**Fig. S1.**
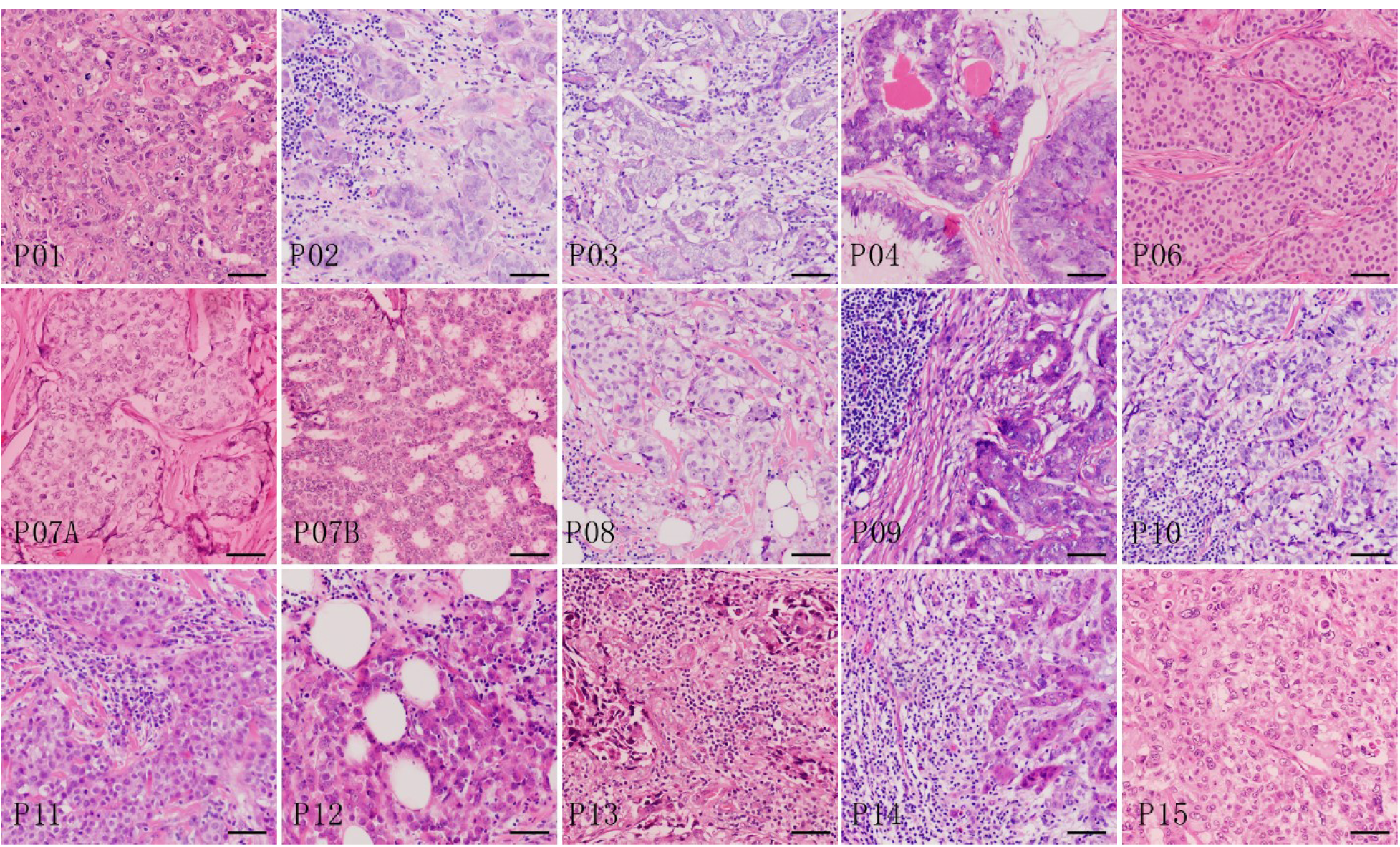
Hematoxylin and eosin staining results of the tumor tissues from each patient. Scale bar, 50μm.

**Fig. S2.**
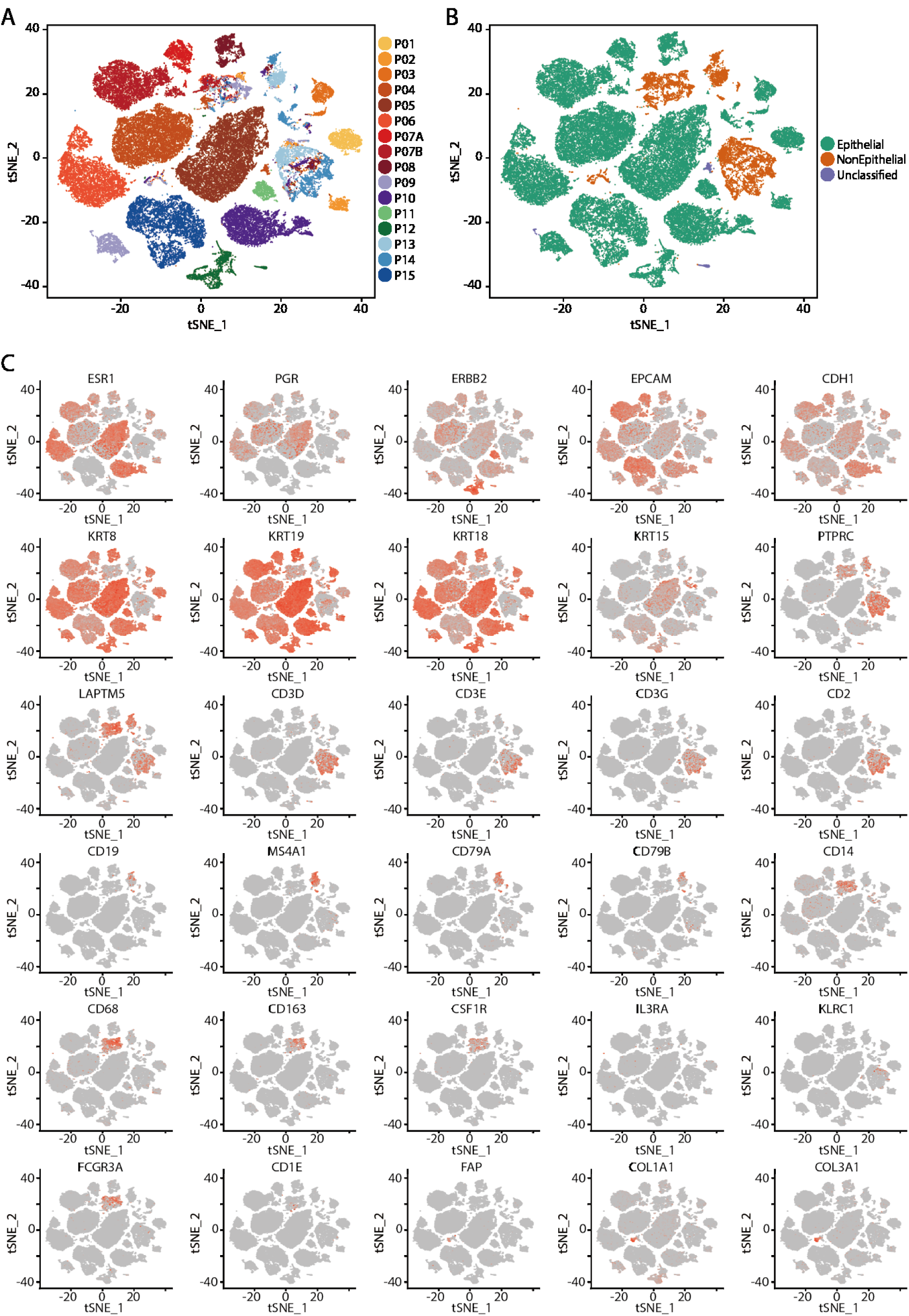
Identification of epithelial cells and non-epithelial cells based on cell markers. (A) t-SNE plot of all cells from 15 patients; the cells are colored according to the sample. (B) The cells are annotated as epithelial cells, non-epithelial cells and unclassified clusters. (C) t-SNE plot of all cells showing the expression of marker genes used for identifying epithelial cells, immune cells and CAFs.

**Fig. S3.**
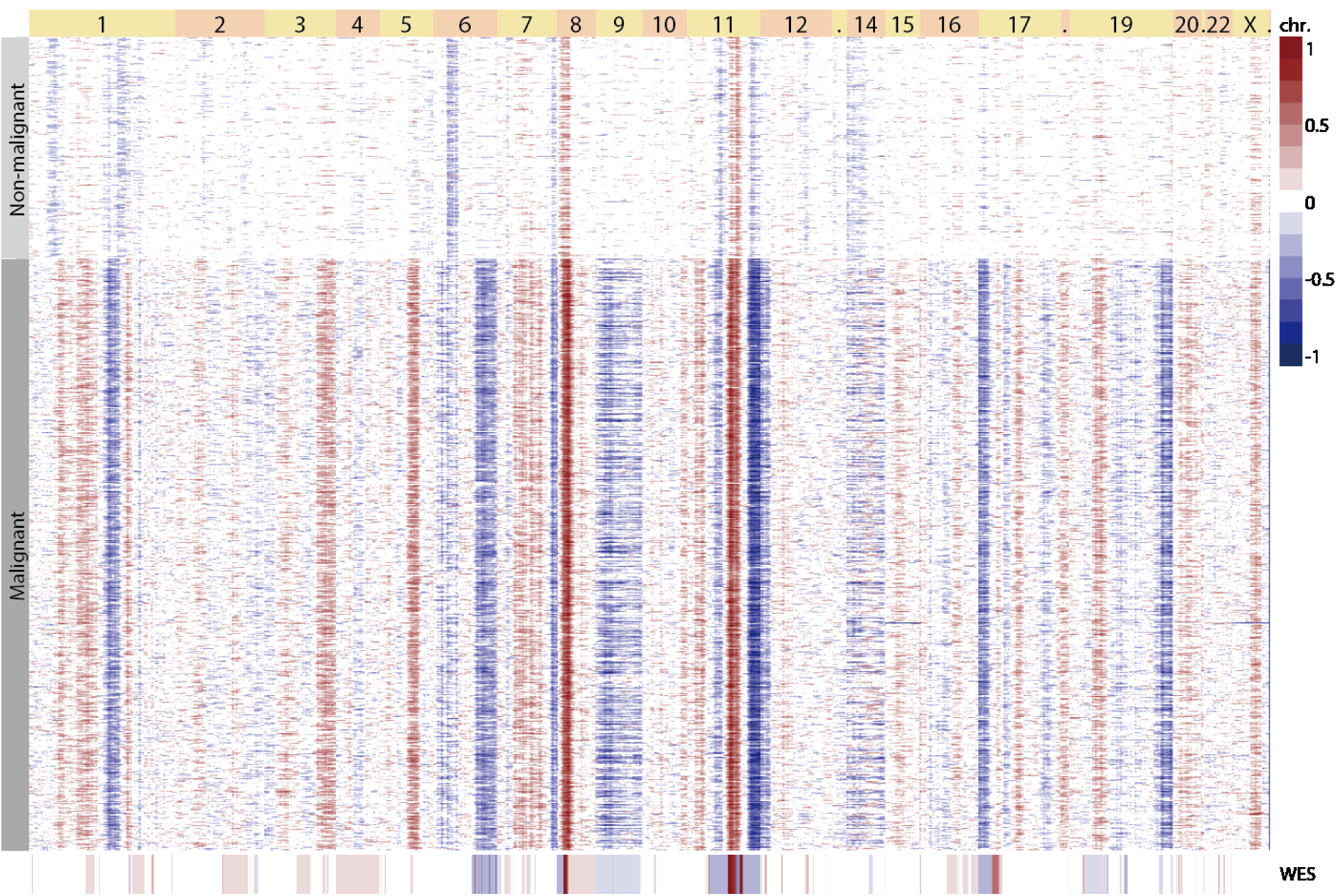
Separate tumor cells from non-tumor cells according to the inferred CNVs. Heatmap showing the inferred CNVs in a representative sample (P03). Red: amplification, blue: deletion.

**Fig. S4.**
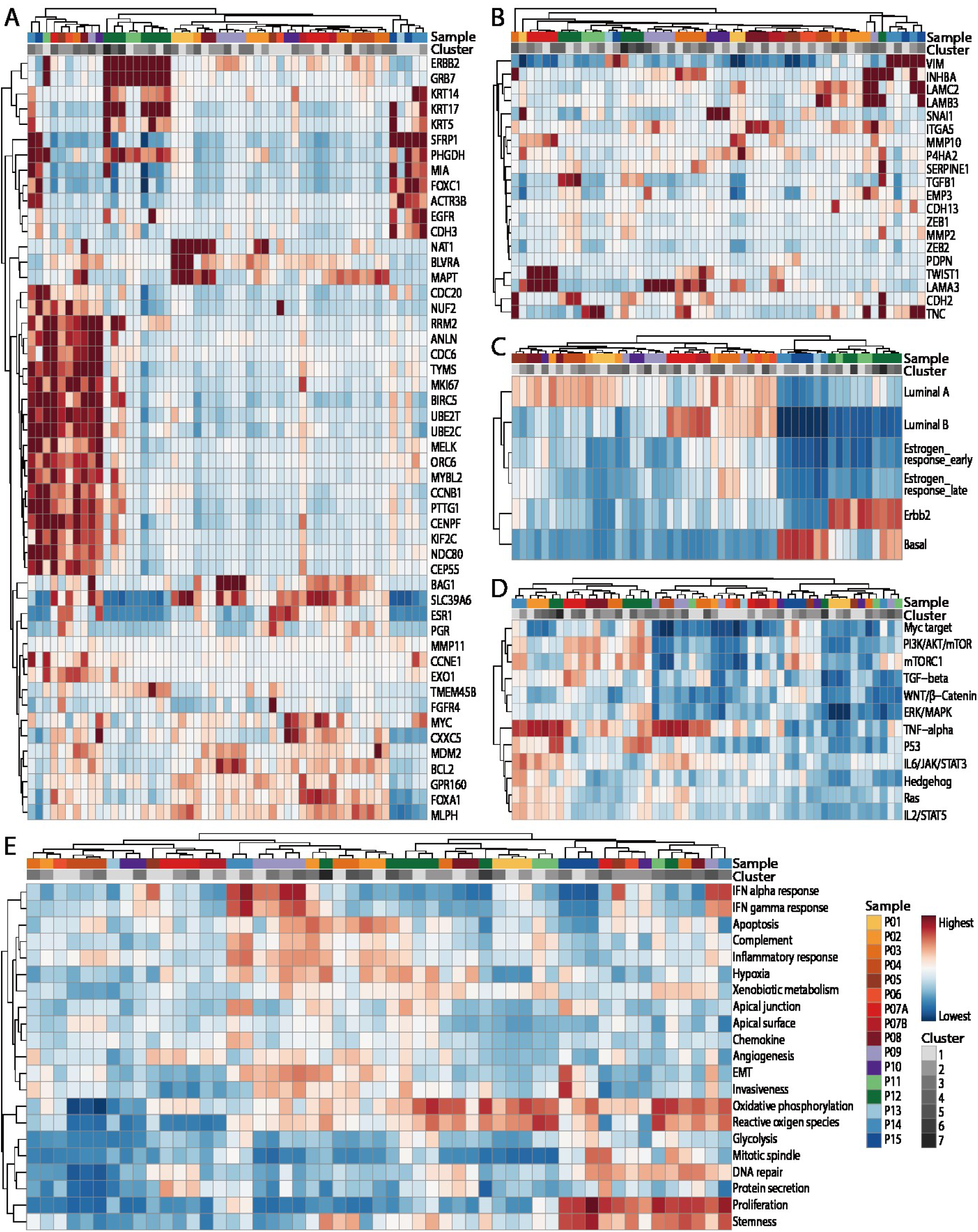
Pathway analysis of the tumor cells across the tumor cell subpopulations. (A) Heatmap showing the average expression of PAM50 genes in each cluster of tumor cells across the 16 samples. (B) Heatmap showing the average expression of EMT-associated genes in each cluster of tumor cells across the 16 samples. (C, D, E) GSVA of breast cancer molecular subtype-associated pathways (C), oncogenic signaling-related pathways (D), and oncogenic phenotype-related pathways in different clusters of tumor cells (E).

**Fig. S5.**
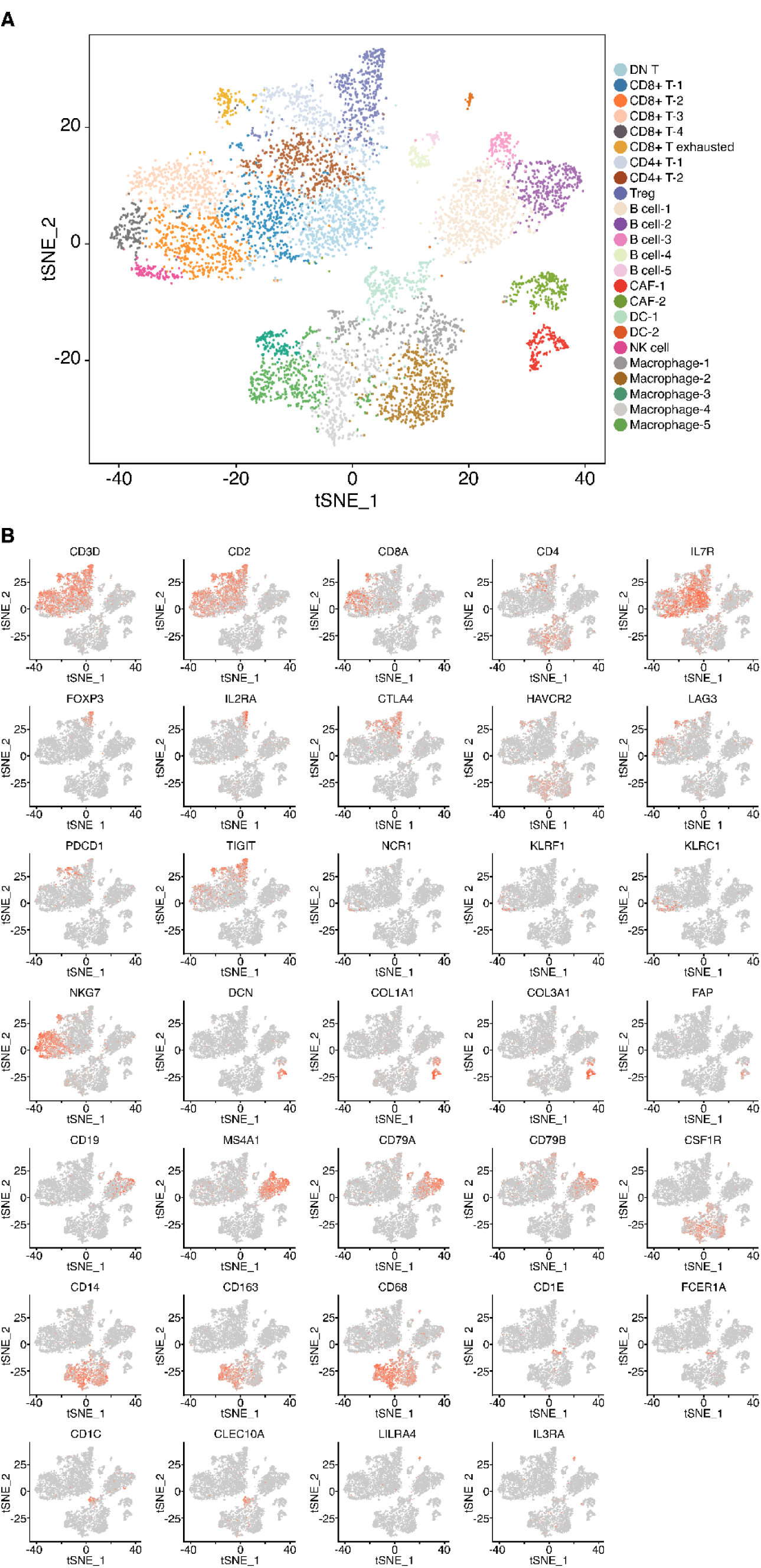
Identification of subtypes of immune cells and CAFs. (A). t-SNE plot of immune cells and CAFs from 15 patients. The cells are colored according to the specific subtype of immune cells and CAFs. (B) t-SNE plot of immune cells and CAFs showing the expression of cell marker genes used for identifying the specific immune cell and CAF subtypes: T cells (CD3D, CD2, CD8A, CD4 and IL7R), Tregs (FOXP3 and IL2RA), exhausted T cells (CTLA4, HAVCR2, LAG3, PDCD1 and TIGIT), NK cells (NCR1, KLRF1, KLRC1 and NKG7), CAFs (DCN, COL1A1, COL3A1 and FAP), B cells (CD19, MS4A1, CD79A and CD79B), macrophages (CD14, CD163 and CD68), dendritic cells (CD1E, FCER1A, CD1C and CLEC10A) and pDCs (LILRA4 and IL3RA).

**Fig. S6.**
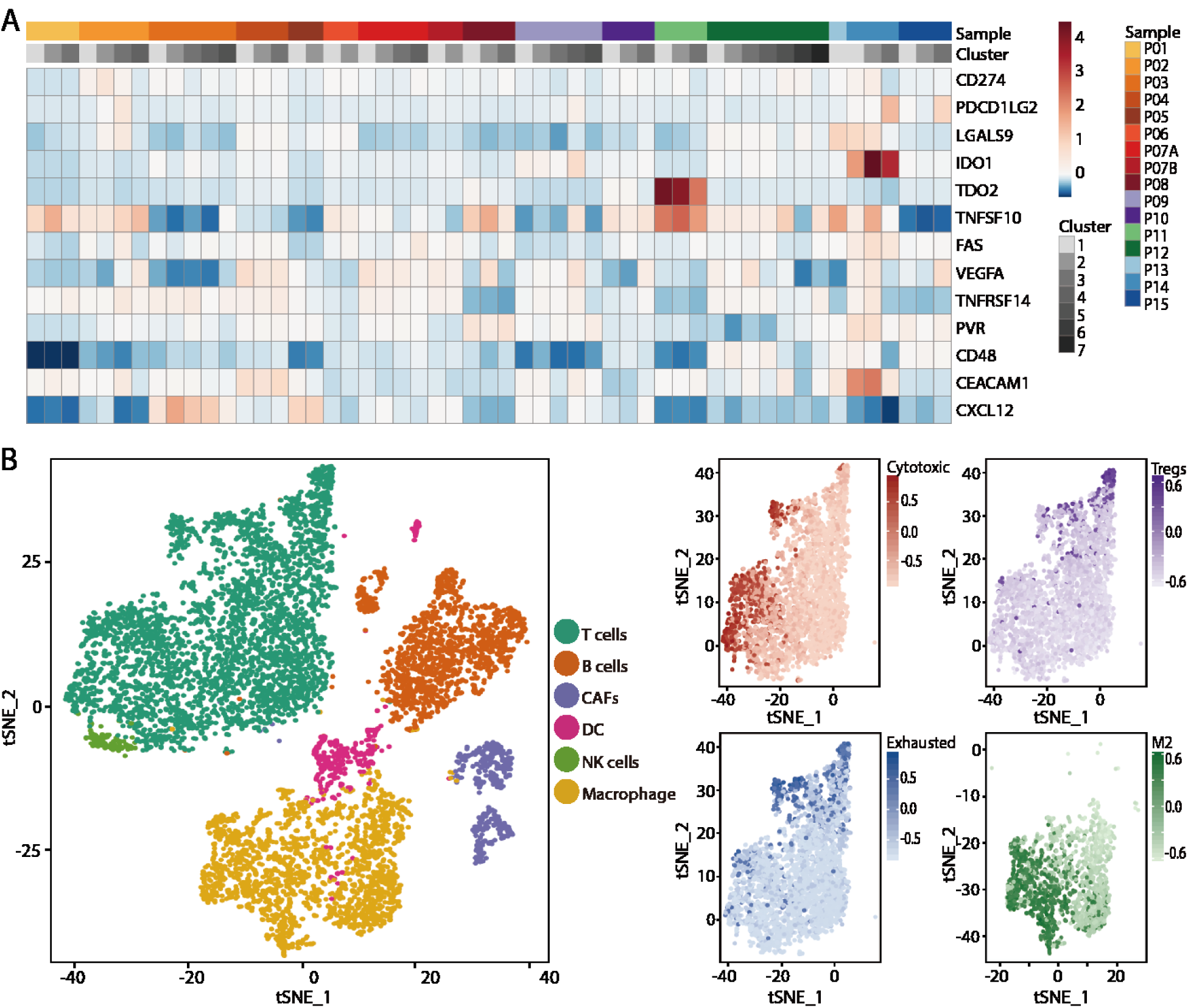
Analysis of the signatures and genes associated with tumor immune evasion. (A) Heatmap showing the average expression of genes in tumor cells associated with immune evasion. (B) t-SNE plot of immune cells and CAFs. t-SNE plot of T cells and macrophages showing the T cell functional states (GSVA scores of cytotoxic, Treg and exhausted gene signatures) and macrophage M2 signatures.

**Fig. S7.**
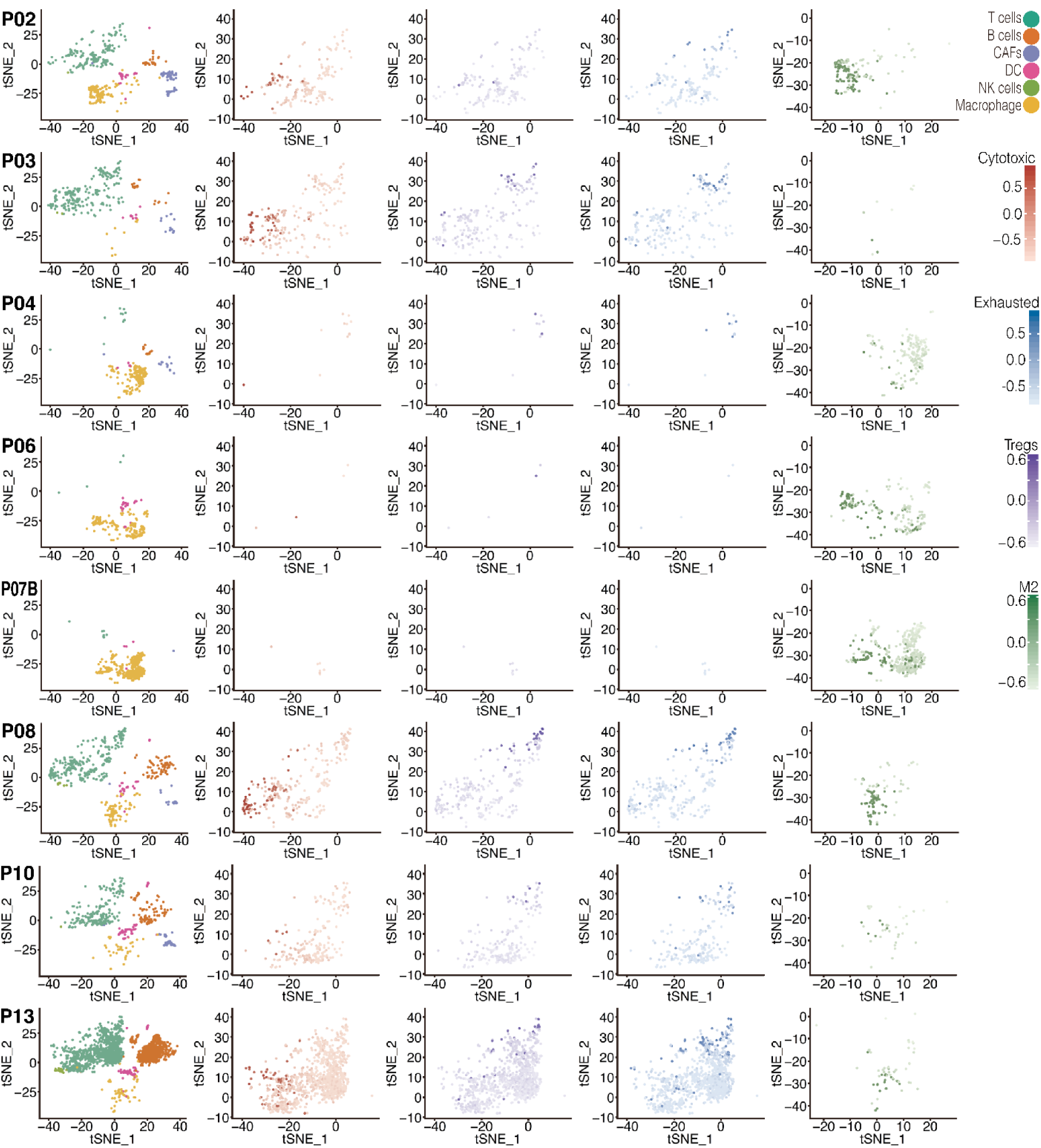
Comprehensive analysis of signatures associated with immune evasion in individual patients. t-SNE plot of immune cells and CAFs in individual patients. t-SNE plot of T cells and macrophages showing T cell functional states (GSVA scores of cytotoxic, Treg and exhausted gene signatures) and macrophage M2 signatures.

**Table S1.**
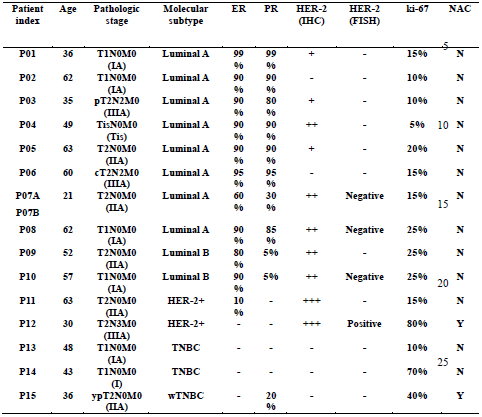
Clinical and histological profiles of the breast cancer specimens. TNBC, triple-negative breast cancer; ER, estrogen receptor; HER2, human epidermal growth factor receptor 2; PR, progesterone receptor; FISH, fluorescence in situ hybridization; Y, neoadjuvant chemotherapy; N, without neoadjuvant chemotherapy.

